# Probabilistic Tracking of U-fibers on the Superficial White Matter Surface

**DOI:** 10.1101/2022.05.05.490829

**Authors:** Xinyu Nie, Yonggang Shi

## Abstract

The short association fibers or U-fibers connect two neighboring gyri and travel in the superficial white matter (SWM) beneath the cortical layer. These U-fibers are essential for the understanding of neurodevelopment and neurodegeneration. However, the complex structures and the high curvature of the U-fibers lead to erroneous streamlines reconstruction of the traditional tractography since the volume-based tractography cannot use the biological characteristic of U-fibers that they tightly beneath the cortical layer. In this work, we proposed a surface-based framework for probabilistic tracking of the U-fibers on the triangular mesh of the SWM. We develop a novel approach to project the fiber orientation distributions (FODs) data onto the tangent space of the SWM surface. With the projected FODs, an advanced probabilistic tracking technique, which regularizes the streamlines based on the intrinsic geometry of the surface, is developed to reconstruct the highly bent U-fibers on the SWM surface. In the experiment, we demonstrate our method based on the high-resolution diffusion imaging data from the Human Connectome Project (HCP). We quantitatively compare the proposed method with state-of-the-art volume-based tractography from MRTrix and another surface-based tractography on the U-fibers of the central sulcus. Moreover, we show the reconstructed U-fibers on the parietal lobe and frontal lobe. The results show that our method outperforms the other two methods and successfully reconstructs the U-fibers on the cortical regions with high intersubject variability.

## 1 Introduction

The short association fiber in the superficial white matter (SWM), which is distributed beneath the cortical surface, is known to be important in the neurodevelopmental during brain maturation [1, 2], in neurodegeneration [3], and in brain pathologies [4-6]. A recent work [7] reported that the U-fibers, assumed as simple wires to connect adjacent gyri, have complex structures and play a vital role in the neural network throughout the entire cerebrum. However, the diffusion-weighted magnetic resonance imaging (dMRI) based technique, tractography is hard to reconstruct the U-fibers [8-10] entirely. Indeed, the complex structures and the high curvature of the U-fibers lead to a false-negative reconstruction of the traditional tractography algorithm [8, 9]. Moreover, the high intersubject variability of the folding patterns leads to the difficulty of clustering and group analysis of the U-fibers [10]. The main difficulty of the traditional tractography is that they are volume-based techniques; they cannot use the essential biological characteristics that the U-fibers travel on the SWM surface.

A surface-based framework [11] tracks the U-fibers on the triangular mesh of the SWM that overcomes the difficulties of volume-based tractography. In this work, they first interpolate the fiber orientation distribution (FOD) [12] data onto the vertices of the preprocessed triangular mesh of the white matter (WM) surface. Then, they pick up the peak directional of the FODs data at each vertex and track the U-fiber on the triangular mesh. For the current point of a track within a triangle, they find a direction for the next tracking point by interpolating and projecting the directional vectors from the vertices of the triangle. This surface-based tracking method outperforms the volume-based tractography on the reconstruction of U-fibers between the precentral and postcentral gyrus. However, this tracking technique is deterministic, thus does not explore all the information of the FOD data. Moreover, the performance of this tracking technique depends highly on the choice of seed points. This method does not work successfully on the cortical regions where the folding patterns are highly variable from subject to subject.

This work develops a probabilistic tracking technique that reconstructs the U-fibers on the SWM surface. Firstly, we develop a method to project the 3D FOD data to the tangent surface of the SWM surface, and the projected 2D FOD data defined in the tangent space can be normalized to a probabilistic distribution. Then, we develop a probabilistic tracking method, which regularizes the tracking by the parallel transport from the differential geometry to reconstruct the U-fibers on the triangular mesh. The proposed tracking technique is intrinsic to the geometry of the surface so that it is better to handle the highly bent U-fibers. The proposed method works on any cortical region and is flexible for the user-defined seed regions.

## 2 Method

### 2.1 FOD Projection

The fiber orientation distribution (FOD) is an advanced model for the diffusion MRI to resolve the crossing fibers problem [12]. For each voxel of the image, the FOD is a symmetric function defined on the unit sphere surface, and the value of the FOD function is a representation of the diffusion strength in the corresponding direction. The FOD data provides an opportunity to reconstruct the deep white matter fiber bundle better. However, since the deep white matter (DWM) crosses with the SWM near the cortical surface, the FOD data represent both the information of the SWM and the DWM. Moreover, the FOD information for the SMW may not lie in the tangent space of the SWM surface due to the partial volume effect in diffusion MRI. To overcome these difficulties, we projected the 3D FOD data to the tangent space of the SWM so that the projected 2D FOD data represent the information of the fibers that travel along the SWM surface. For each point on the surface of the SWM, we use the following formula (1) to project the 3D FOD function to the tangent space of the SMW.

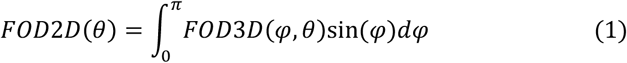

Where the reference plane for the spherical coordinate system is the tangent plane of the SWM surface, and the zenith direction is the normal direction of the tangent plane. A local spherical coordinate system on the triangular mesh is shown in Figure1. (a), an edge e_1_ of the triangle is chosen as the axis whose azimuthal angle θ equals 0, and the polar angle φ is the angle to the zenith direction.

**Figure 1.**
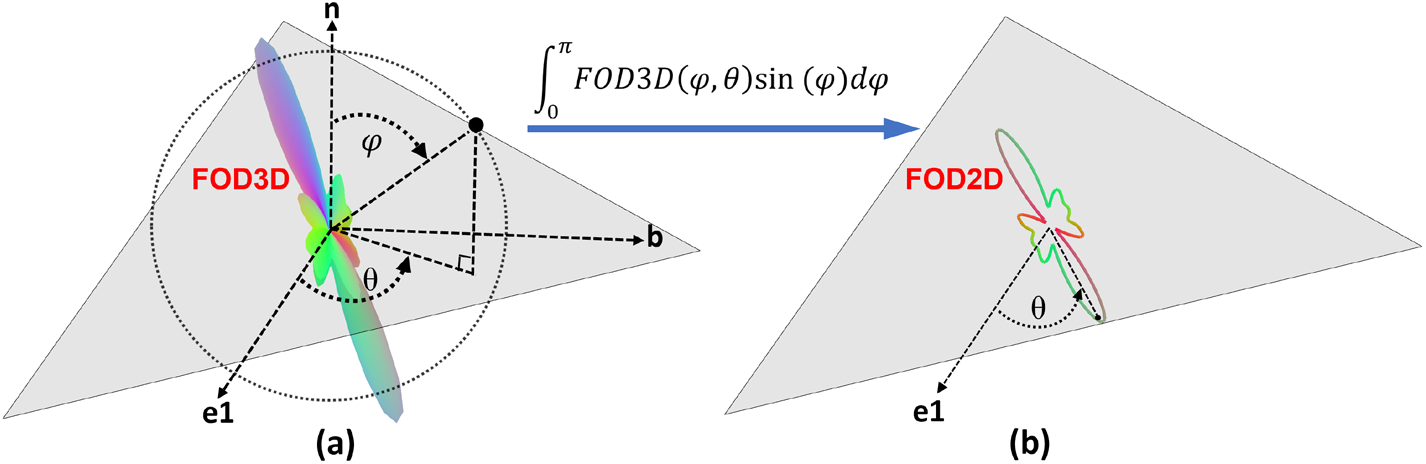
(a) The FOD3D function is represented in the local spherical coordinate system defined on the center of a triangle of the SWM mesh. The reference plane of the system is the triangle plane, mean-while the zenith direction aligned with the normal direction n. The axis with 0 azimuthal angle θ is aligned with an edge e1 of the triangle. (b) The diffusion information is projected onto the triangle plane using formula (1). The FOD2D function is parametrized by θ in the local coordinate system and normalized to be a probability distribution function on the unit circle S^1^.

The weighted term sin(φ) in the integral eliminates the diffusion information that is perpendicular to the SWM surface while reserving the information that travels along the surface. The projected 2D FOD function parametrized by the azimuthal angle θ in the local coordinate system is defined on a unit circle S^1^ which lies in the tangent space of the SWM surface as in Figure1. (b). The value of the FOD2D function represents the diffusion strength along the corresponding direction on the surface. Numerically, for each triangle of the triangular mesh, we use the spline interpolation to interpolate the FODs from neighboring voxels to the center point. We now can compute the FOD3D on the spherical coordinate system at the center point of the triangle using the spherical harmonics (SPHARM) model of the FODs. Then the FOD2D is computed using formula (1). The FOD2Ds are piecewise constant function fields on the triangular mesh; namely, each FOD2D function is defined on a triangle plane and represents the diffusion information within this triangle. Figure1 shows the FOD3D function on a triangle of the triangular mesh and then is projected onto the triangle plane.

### 2.2 Tracking on the Triangular Mesh

The reject sampling method is used to sample the direction of the streamline based on the normalized FOD2D function, which is a probability distribution function on the unit circle S^1^. Given a seed triangle, we choose the center point of the triangle as the seed point, and we randomly sample a direction at this seed point for tracking. Since the FOD2D function is symmetric concerning the center and there are symmetric directions to track on the seed point, we track the opposite directions respectively and combine them to a complete streamline.

#### Tracking within a Triangle

For a current tracking point within a triangle (including the boundary points) with a given direction, we compute the straight-line segment, which is aligned with the given direction, originates from the current point, and terminated at the triangle boundary. Then the terminated point becomes the new current tracking point and enters into a new adjacent triangle. Since our FOD2D function does not change within a triangle (like a piecewise constant vector field), the track is a straight line within each triangle once the direction is accepted. Thus, our tracking points always lie in the boundary of the triangles except for the seed point.

#### Tracking across two Triangles

When our current point leaves a previous triangle and enters into a new adjacent triangle, we have to search for a new direction for further tracking. Like the volume-based tractography, the new direction is determined by both the sampling function and some regularizations (e.g., angle difference, curvature). However, the regularizations used in volume-based tractography are not helpful for surface tracking since the SWM surface, although embedded in Euclidean space R^3^, has its intrinsic geometry. The challenging endeavor is how to compare two nearby tangent vectors, namely, how to regularize the sampling direction in the new triangle according to the direction vector in the previous triangle. An affine connection (or covariant derivative) introduces such a geometric structure on a manifold to compare vectors in neighboring tangent spaces [13]. Since the SWM mesh embedding in R^3^ and inherited Riemannian metric from R^3^, the affine connection (Levi-Civita connection) induced by the Riemannian metric is natural to be used. With the Levi-Civita connection, a vector in a tangent space can be parallel transported (the covariant derivative along the path is 0) to another tangent space. We can parallel transport the previous direction vector and compare it with the sampling direction vectors in the new triangle. The parallel transport induced by the Levi-Civita connection is simple for the adjacent triangles on the triangular mesh (Figure2); namely, the parallel transport is achieved by unfolding the pair of adjacent triangles, translating the vector from one triangle to the other since the two triangles are now in the same plane, and finally folding the triangles back to original position [14]. Since we use a local coordinate system for each triangle plane as discussed in the FOD projection, we must transfer the vector from the local system to the R^3^ for parallel transportation. For a direction parametrized by θ in the local coordinate system of a triangle T_1_ as in Figure2. (a), the corresponding vector v in R^3^ is acquired by

**Figure 2.**
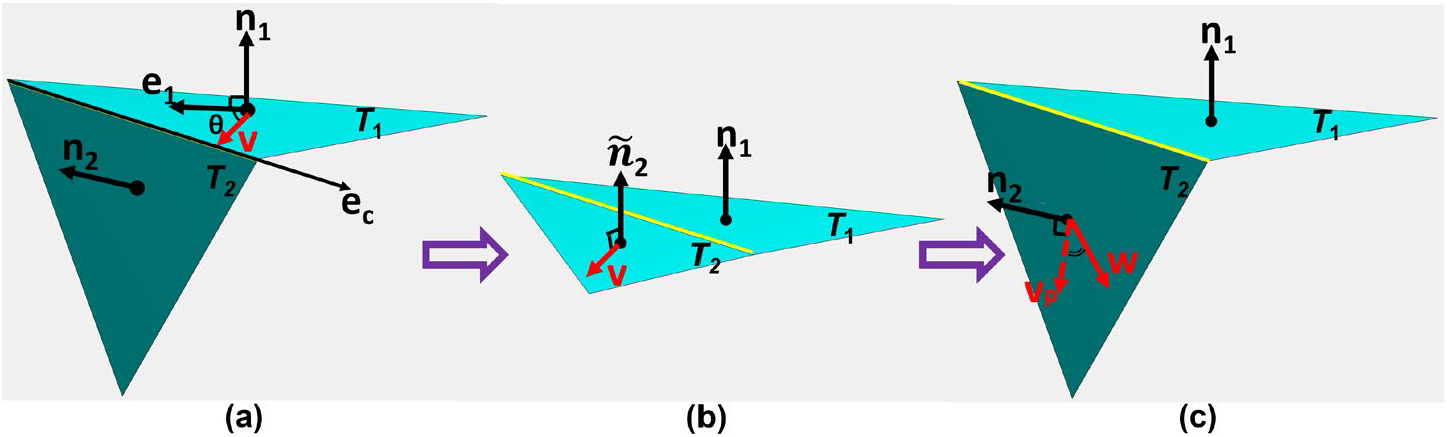
Parallel Transport: The parallel transport consists of unfolding a pair of triangles to a common plane (a)∼(b), translating the red tangent vector v from triangle T1 to T2 (b), and then folding back the triangles to the original position (b)∼(c), the transported vector vp can be compared with the sampling direction w in the triangle T2.

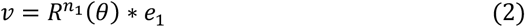

Where e_1_ and n_1_ are the directional vectors of the axis edge and normal as in Figure2, and the matrix R is the operator that rotates by angle θ around the normal direction. The parallel transport from triangle T_1_ into T_2_ is achieved by rotating the vector v

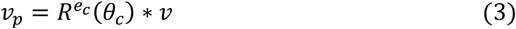

Where e_c_ is the direction vector of the common edge of the two adjacent triangles, and the angle θ_c_ is the angle between the normal vectors of the two triangles (The two triangles must be oriented in the same way). The vector v_p_ lies in the space of the triangle T_2_ and can be compared with the new sampling direction in triangle T_2_ as in Figure2. (c). We repeat sampling by FOD2D at the new triangle until the angle difference between the sampled direction and the parallel transported direction v_p_ is less than a threshold θ_in_. The accepted sampling direction is used to track within the new triangle using the algorithm discussed in the last paragraph.

For the extreme case that the tracking leaves the previous triangle at a vertex (hardly ever happens in practical application), we use the geodesic polar map [15] to flatten the one ring of the vertex into a 2D chart. In the 2D chart, we easily find the upcoming triangle since we can extend the tracking line through the vertex. Finally, we map the extended tracking line from the 2D chart to the original upcoming triangle in the 3D mesh by the inverse geodesic polar map, and the direction vector of the mapped line is used to regularize the sampling as before.

We repeat tracking within the triangle and across the triangles until the tracking point encounters the boundary of the ROI or the FOD2D data fails to generate a valid direction. Finally, the two tracks with opposite directions and originating from the same seed point are combined to a complete streamline.

Since we use parallel transport to regularize tracking, the tracking method is intrinsic to the surface. Our tracking method works well in the cortical regions where the surface bends dramatically, and the volume-based tractography often fails. Also, our tracking technique does not depend on the choice of the seed points; the user can choose the seed regions on the cortical surface wherever he likes. For the U-fibers that connect two neighboring gyri, we recommend the user select seeds in the sulcal areas and choose the gyral regions as the stopping criteria. Our algorithm takes about 1-2 hours to generate around 3000 streamlines for U-fibers of the central sulcus on a laptop with a 2.6-GHz Intel CPU. Our code is compatible with parallel computation for faster computation.

## 3 Experiment

We apply the surface-based probabilistic tracking method to the large-scale 484 subjects from the Human Connectome Project (HCP) [16]. We reconstruct fiber orientation distributions (FODs) using the method [12] from the multi-shell diffusion MRI data. From the T1-weighted MRI data, the triangular mesh representation and parcellation of the gray matter (GM) and white matter (WM) of the cortical surface is processed by the FreeSurfer [17, 18]. The WM mesh is deformed inward toward the DWM along the normal direction by half of the voxel size to generate the SWM mesh surface for the tracking of U-fibers.

In the experiment, we compare the proposed method with state-of-the-art MRtrix software [19] that is the conventional volume-based tractography, as well as the surface-based deterministic framework [11]. We compare the three methods of reconstructed U-fibers in the precentral and postcentral cortex, where U-fibers are shown in several kinds of literature [20-22]. The gyral skeletons of the GM surfaces are computed using the graph-cut-based method [23]; meanwhile, the central sulcus was detected within the 2mm-geodesic distance transforms from the precentral and postcentral ROIs, respectively. All experiments are on the left hemisphere.

For the surface-based deterministic method, to follow the parameters in work [11], we set up the step size to be 0.1mm, and the angle thresholds as TH_θin_ = 1 degree and TH_θxing_ = 10 degrees. For the MRtrix, the IFOD1 algorithm is used; we set up the step size to be 0.1mm and the maximum angle of steps as 10 degrees. For the proposed surface-based probabilistic method, we set up the angle thresholds θ_in_ as 10 degrees to match other methods. Also, we set the minimum and maximum length of any streamline of any method to be 20 and 80mm for the short association fibers as used in [24]. We choose the sulcal area on SWM mesh as the seed region for the two surface-based algorithms and extend the sulcal seed region for the volume-based method to the neighboring voxels within 6mm. We randomly sampled seed points in the seed area until 3000 streamlines were generated for each method. The reconstructed U-fibers of two subjects for all methods are shown in Figure3.

**Figure 3.**
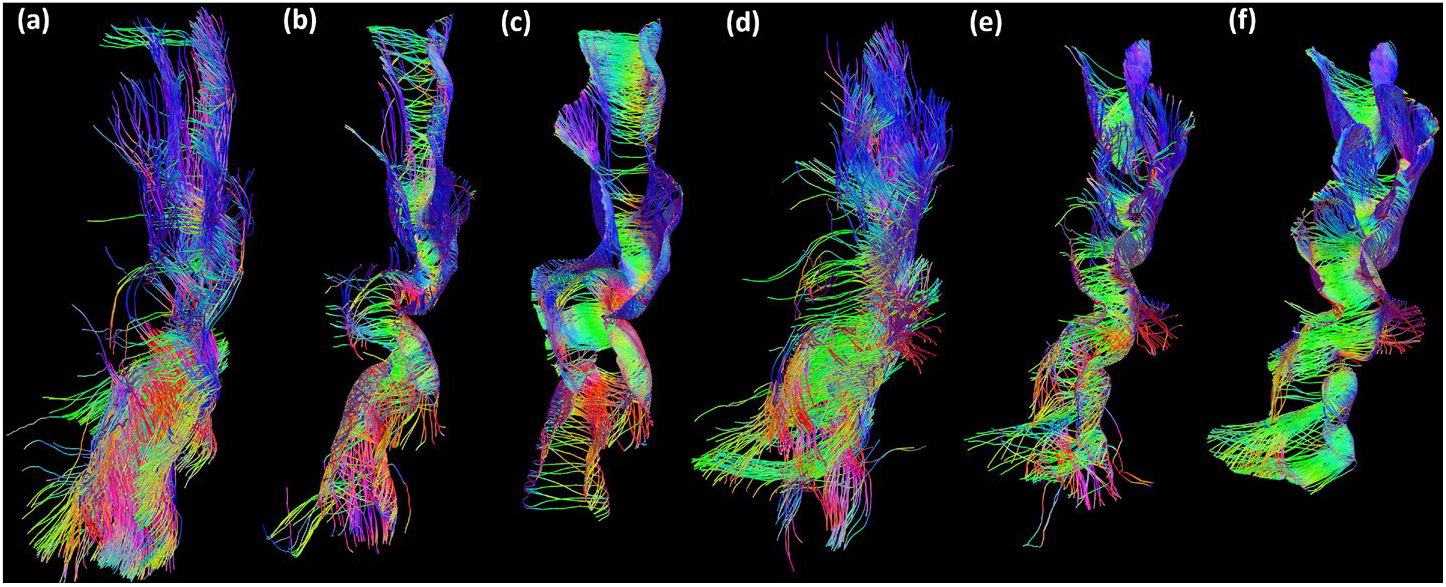
The U-fibers reconstruction of the central sulcus for subject 101107 (a, b, c) and subject 106016 (d, e, f) by three methods. MRTrix (a, d), the surface-based deterministic algorithm (b, e), and the surface-based probabilistic algorithm (c, f). We down-sample the number of streamlines to be 1000 for each method.

We use three citeria for quantitative evaluation of the U-fiber: ‘well-distributed’, ‘well-U-shaped’, and the Topographic Regularity. The ‘well-U-shaped’ is not a strict evaluation for the quality of the U-fiber but to show that the surface-based methods can generate streamlines with higher curvature. We first average the pointwise curvature for an individual streamline, then compute the mean curvature for all the streamlines within a subject. Figure4. (a) shows the mean curvature for all the 484 subjects. To measure ‘well-distributed’, we uniformly partitioned both the precentral and postcentral gyral skeletons into 20 sections and counted how many sections were hit by the streamlines, namely, one section is hit if the endpoint of a streamline is within 3mm to the section. Figure4. (c)∼(d) show the ‘well-distributed’ measure of all three methods, and each boxplot shows the number of the sections hit by the streamlines for the 484 subjects. The topographic regularity is an important property widely presented by the fiber bundles [25-28]. We measure the topographic regularity of the U-fiber using the most intuitive metric proposed in the work [25]. For each subject, we use the classical multidimensional scaling (MDS) to project both the beginning points (precentral gyrus) and the endpoints (postcentral gyrus) of the streamlines to the 2D plane, respectively. Then we compute the Procrustes distance to measure the shape difference between the beginning points and endpoints in the 2D plane. The Procrustes distance of the U-fibers for all the 484 subjects is shown in Figure4. (b).

**Figure 4.**
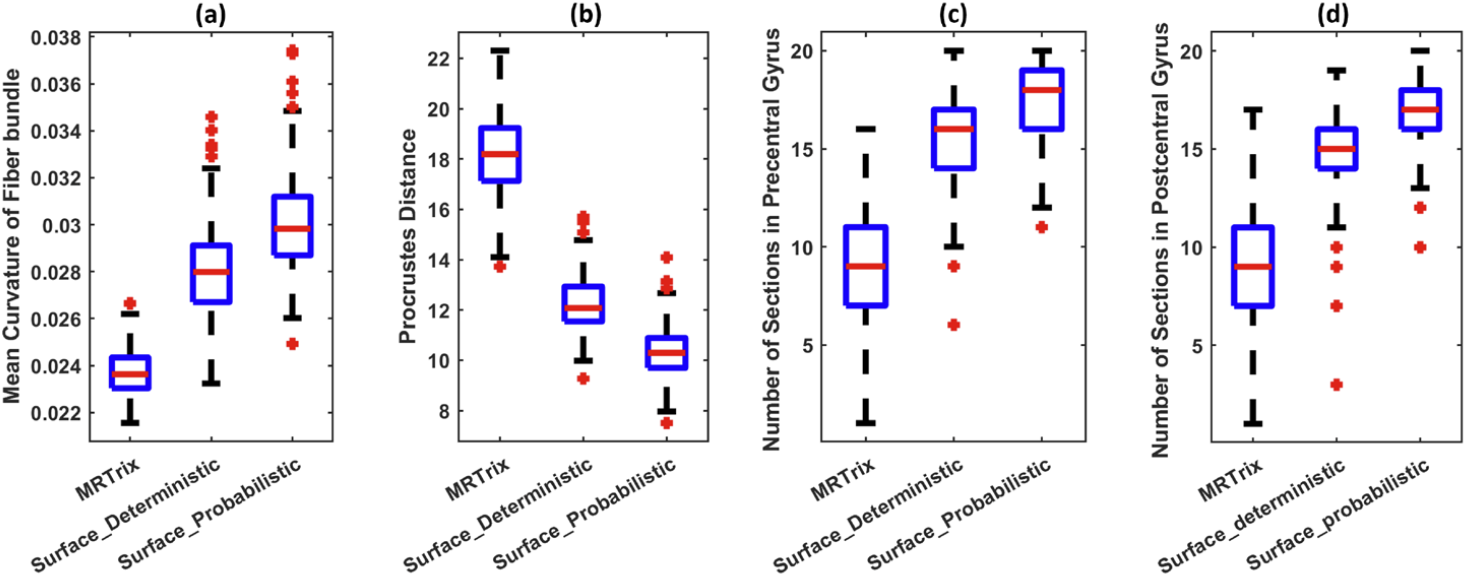
Quantitative evaluation of the three methods using the large-scale datasets of 484 HCP subjects. (a) Box plots the ‘well-U-shaped’, we compute the mean curvature of all the streamlines for each subject. (b) Box plots of the Procrustes distance to show the Topographic regularity of the U-fibers. (c)∼(d) Box plots of the ‘well-distributed’ measure, namely, the number of the gyral sections hit by the streamlines from each method

The ‘well-distributed’ measurements show that the U-fibers reconstructed by the proposed method cover more gyrus regions. Moreover, the lower Procrustes distance shows that U-fibers provided by the proposed method are more topographically regular, which implies the streamlines are well-organized; meanwhile, the mean curvature results show that the proposed method can generate more bent U-fibers. Two examples in Figure3 give some explanations for the quantitative results. Most streamlines reconstructed by the volume-based tractography do not closely follow the cortical surface, while both surface-based methods generate streamlines that follow tightly to the cortical surface. The proposed surface-based probabilistic method provides similar but superior-quality U-fibers compared with the surface-based deterministic method since the probabilistic tracking technique explores more diffusion information while regularizing the track by the intrinsic geometry of the SWM surface.

We also apply the proposed method to more challenging cortical regions with the same parameters used in the central sulcus. Figure5 shows the reconstructed well-organized U-fibers that connect the inferior and superior parietal lobules; even the cortical folding patterns around the intraparietal sulcus are highly variable. Moreover, the reconstructed U-fibers on the whole frontal lobe are shown in Figure6 to demonstrate that our method can reconstruct the U-fibers on a wide range of cortical areas.

**Figure 5.**
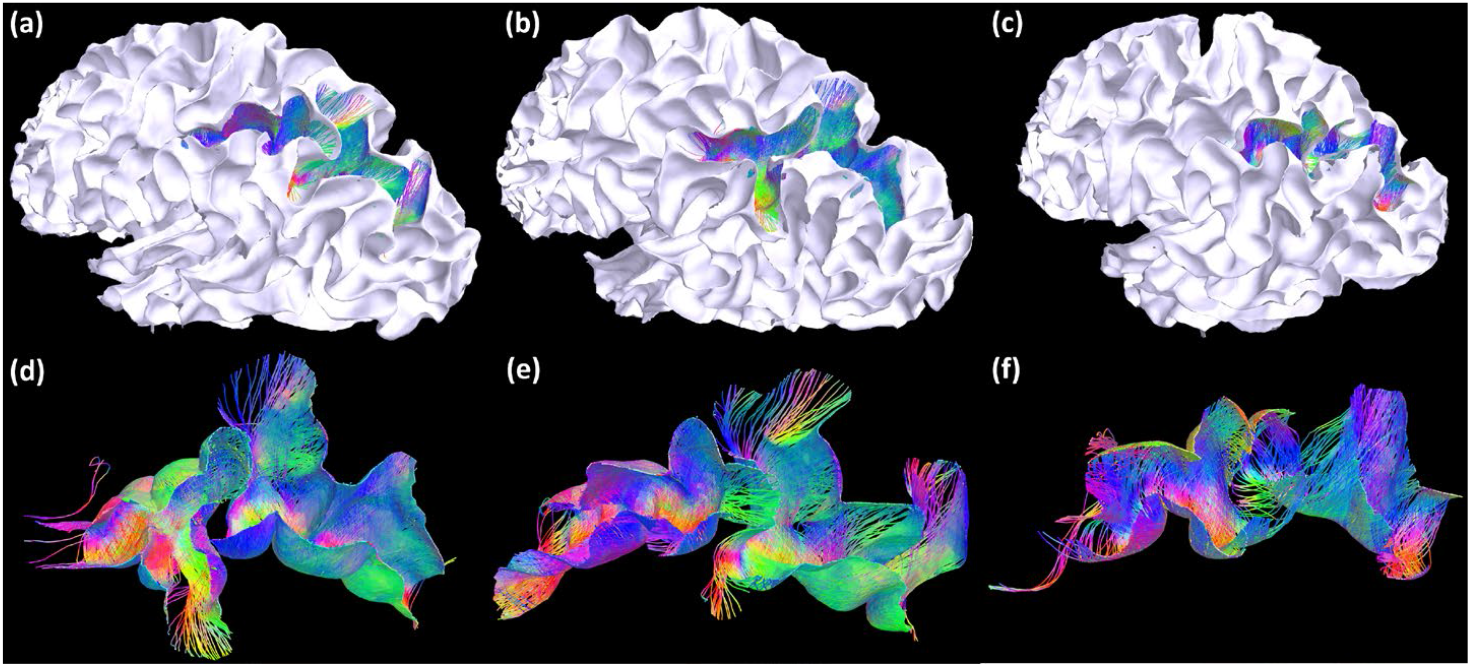
The reconstructed U-fibers that connect the inferior and superior parietal lobules for subject 106016 (a, d), subject 101915 (b, e), and subject 108121 (c, f). The streamlines are superimposed on the SWM surface in (a, b, c). The same streamlines are plotted without the surface in (d, e, f).

**Figure 6.**
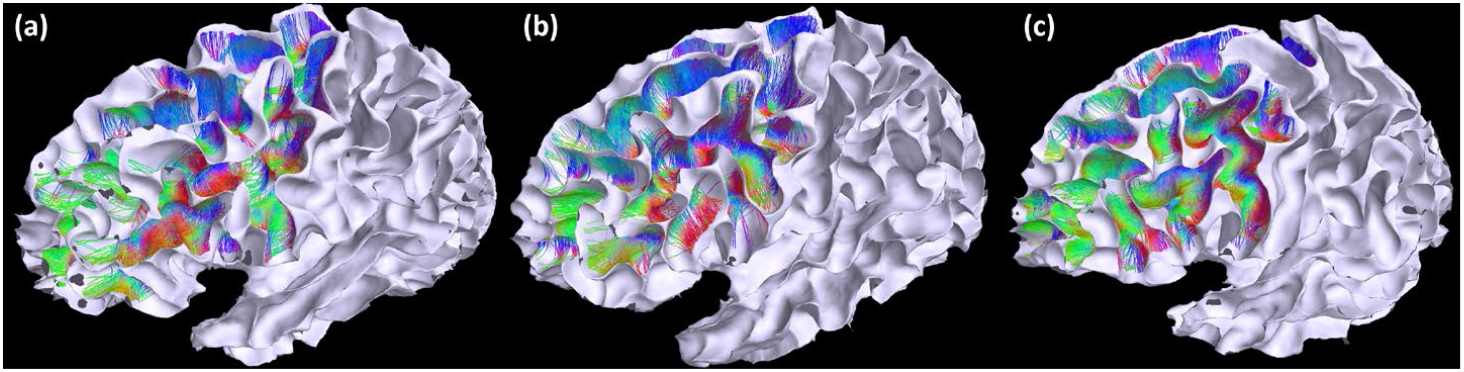
The reconstructed U-fibers on the frontal lobe: (a) subject 106016, (b) subject 101915, and (c) subject 108121. The streamlines are superimposed on the SWM surface.

## 4 Conclusion

This paper proposes a novel surface-based probabilistic framework to track the U-fibers on the SWM surface. Our method projects the FOD information onto the SWM surface and tracks the U-fibers using the intrinsic geometry of the surface. The experiments show that our method outperformed the other two tracking methods by providing U-fibers with higher quality. We will reconstruct the U-fibers on more cortical regions and apply the proposed algorithm to more clinical data in future work.

## Acknowledgement

This work was supported in part by the National Institute of Health (NIH) under Grant RF1AG056573, Grant RF1AG064584, Grant R01EB022744, Grant R21AG064776, Grant R01AG062007, Grant P41EB015922, and Grant P30AG066530; in part by the Human Connectome Project, WU-Minn Consortium (Principal Investigators: David Van Essen and Kamil Ugurbil) funded by the 16 NIH Institutes and Centers that support the NIH Blueprint for Neuroscience Research under Grant 1U54MH091657; and in part by the McDonnell Center for Systems Neuroscience, Washington University. (Corresponding author: Yonggang Shi.)

